# Exponentially decreasing exposure of antigen generates anti-inflammatory T-cell responses

**DOI:** 10.1101/2023.09.15.558014

**Authors:** Arezoo Esrafili, Joshua Kupfer, Abhirami Thumsi, Madhan Mohan Chandra Sekhar Jaggarapu, Abhirami P. Suresh, Aleksandr Talitckii, Taravat Khodaei, Srivatsan J. Swaminathan, Shivani Mantri, Matthew M Peet, Abhinav P. Acharya

## Abstract

Rheumatoid Arthritis (RA) is a chronic debilitating disease characterized by auto-immune reaction towards self-antigen such as collagen type II. In this study, we investigated the impact of exponentially decreasing levels of antigen exposure on pro-inflammatory T cell responses in the collagen-induced arthritis (CIA) mouse model. Using a controlled delivery experimental approach, we manipulated the collagen type II (CII) antigen concentration presented to the immune system. We observed that exponentially decreasing levels of antigen generated reduced pro-inflammatory T cell responses in secondary lymphoid organs in mice suffering from RA. Specifically, untreated mice exhibited robust pro-inflammatory T cell activation and increased paw inflammation, whereas, mice exposed to exponentially decreasing concentrations of CII demonstrated significantly reduced pro-inflammatory T cell responses, exhibited lower levels of paw inflammation, and decreased arthritis scores in right rear paw. The data also demonstrate that the decreasing antigen levels promoted the induction of regulatory T cells (Tregs), which play a crucial role in maintaining immune tolerance and suppressing excessive inflammatory responses. Our findings highlight the importance of antigen concentration in modulating pro-inflammatory T cell responses in the CIA model. These results provide valuable insights into the potential therapeutic strategies that target antigen presentation to regulate immune responses and mitigate inflammation in rheumatoid arthritis and other autoimmune diseases. Further investigations are warranted to elucidate the specific mechanisms underlying the antigen concentration-dependent modulation of T cell responses and to explore the translational potential of this approach for the development of novel therapeutic interventions in autoimmune disorders.

## Introduction

Vaccines often rely on sustained release of antigen from the injection site. For example, subunit vaccines injected with alum are sustainably released to generate a pro-inflammatory response [1,2]. Moreover, vaccines such as DNA and RNA based vaccines infect the host cells and allow for increasing level of antigen availability for processing and display by antigen presenting cells [3,4]. Most of the vaccines are targeted toward increased pro-inflammatory responses and therefore, adjuvants are added to the vaccines to induce antigen presenting cells to generate an antigen-specific adaptive pro-inflammatory B-cell and T-cell response. In the absence of adjuvants, however, a lower quality and quantity of response is generated against the antigen [5]. Interestingly, in the absence of an adjuvant, the effect of rate of exposure of antigen to the immune system is not well understood. For example, during infection the rate at which the antigens are presented to the adaptive immune cells exponentially increases. On the other hand, when the pathogens are cleared these antigens displayed by the antigen presenting cells might decrease in quantity as time progresses. To mimic and study this process, controlled release systems can be utilized.

Several controlled release systems have been developed to release immune-active molecules to modulate immune responses. For example, poly(lactic-co-glycolic) acid polymer-based micro -and nanoparticles have been developed for delivery of adjuvants to generate vaccine-like response [6–10]. Furthermore, it has also been demonstrated that controlled release of vaccines and different dosing regimens provide differential immune responses [11–14]. These studies have mostly focused on generating pro-inflammatory responses by delivering adjuvant and antigen simultaneously and have not studied the effect of antigen delivery only and testing the immune responses to decreasing antigen levels over time.

In an autoimmune disease such as Rheumatoid Arthritis (RA), antigens processed and displayed to the adaptive immune response might be increasing, as the disease progresses. This is observed in clinics where the disease generally starts in peripheral joints such as fingers and then spreads throughout the body, potentially due to increased expression of collagen as the antigen over time. Thus, reducing the display of collagen antigen might be a strategy to decrease inflammation in RA. However, the effect of changing or decreasing levels of collagen on immune responses in the context of RA is not known.

In this study we developed a controlled release system based on Poly(lactic-co-glycolic) acid – Polyethylene glycol – Poly(lactic-co-glycolic) acid (PPP), to deliver antigens (ovalbumin and bovine collagen type 2) in a sustained decreasing manner. The effect of these formulations on T cell responses were then studied *in vivo* in a collagen induced arthritis (CIA) mouse model of RA, which is a widely utilized model of human RA.

## Results

### PPP release OVA in a sustained and exponentially decreasing trend

To generate an exponentially decreasing level of OVA release, biomaterials were utilized. Specifically, PPP dissolved in PBS were admixed with OVA resuspended in PBS at 4 ºC. It was assumed that 100% of OVA was dispersed within the PPP matrix. This solution was then brought to 37 ºC to generate a gel **(Figure 1a)**, and release kinetics were performed in PBS. It was determined that different concentration of PPP in PBS released OVA differentially, with 100 mg/mL of PPP releasing OVA the fastest **(Figure 1b)**. Hence, in order to investigate the impact of exponentially decreasing levels of OVA on immune responses, we employed the slowest release profile achieved through the use of 200 mg/mL PPP for subsequent experiments.

**Figure 1:**
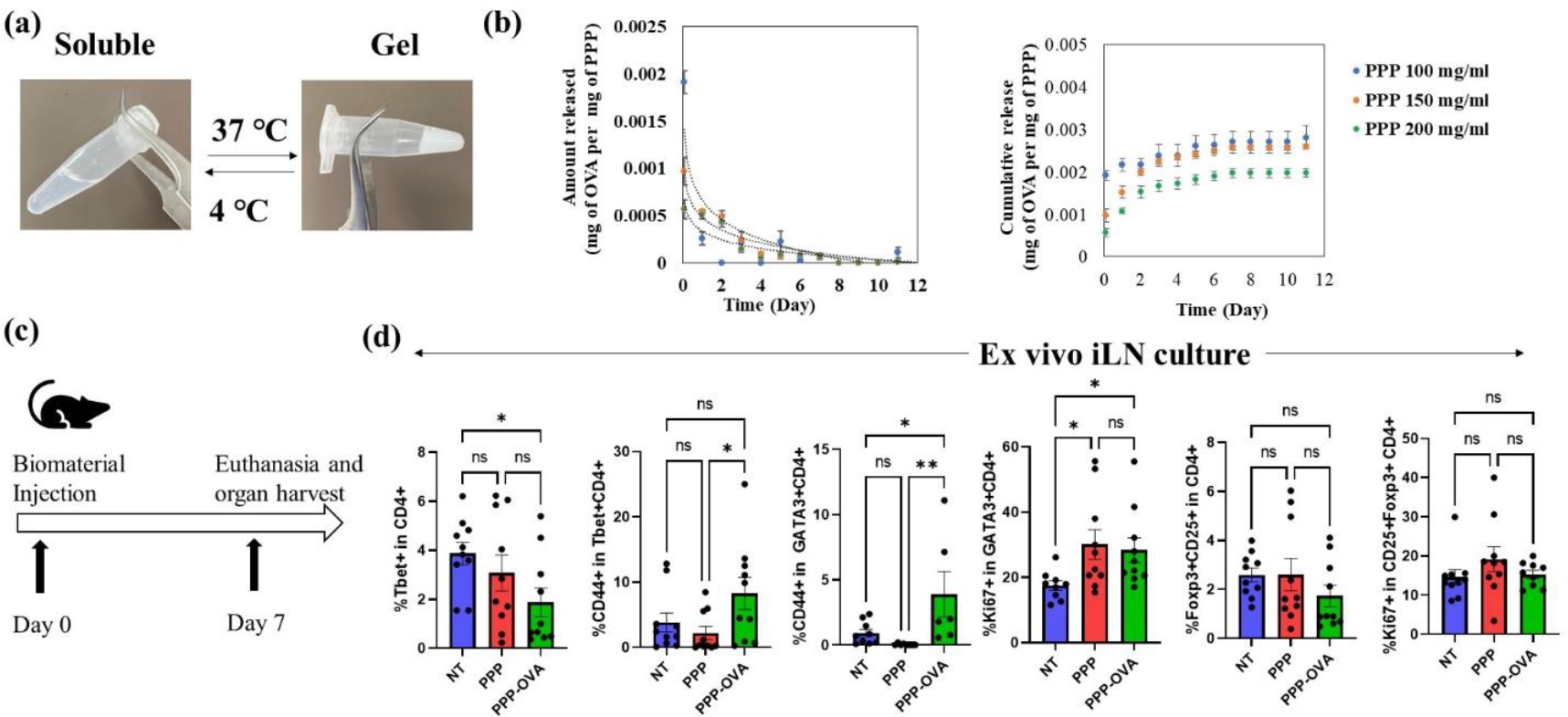
Exponentially decreasing release of ovalbumin (OVA) modulates T cell responses in mice. **(a)** PPP are thermoresponsive in nature, and gel at 37 ºC. **(b)** OVA is released in an exponentially decreasing fashion from PPP, with 200 mg/mL PPP showing slowest release kinetics. **(c)** Schematic of animal experiment to investigate effect of exponentially decreasing release of OVA on T cells. **(d)** Ex vivo culture responses to exponentially decreasing OVA is shown for T helper type 1 (Tbet+CD4+), activated T helper type 2 (CD44+GATA3+CD4+), proliferating T helper type 2 (Ki67+GATA3+CD4+), activated T helper type 1 (CD44+Tbet+CD4+), regulatory T cells (CD4+CD25+Foxp3+), proliferating regulatory T cells (CD4+CD25+Foxp3+). N = 5 mice per group, n = 5-10 technical replicates, Avg±SE, One-way ANOVA with Fisher’s LSD test, * p-value = 0.05 – 0.01, ** -p-value < 0.01, *** - p-value < 0.001.

### PPP releasing OVA with a decreasing trend modulate T cell responses in wild type mice

In this study, we aimed to investigate the effect of PPP releasing ovalbumin (OVA) with a decreasing trend, on T cell responses in a wild-type C57BL/6j mouse model. To achieve this, PPP as a delivery system for OVA was subcutaneously injected on day 0, allowing for sustained release with a decreasing concentration over time, with PPP without OVA (biomaterial control) and PBS (vehicle control) were used as controls **(Figure 1c)**. The rationale behind this approach was to mimic a natural antigen exposure pattern, which can influence T cell responses. By modulating the antigen presentation dynamics, we hypothesized that T cell responses would be affected as well. Therefore, on day 7, mice injected with biomaterials were euthanized and inguinal lymph nodes were isolated. T cells isolated from the inguinal lymph node of the treated mice were then cultured in the presence of OVA or bovine serum albumin (BSA, antigen control) to assess how T cells respond in the presence of antigen. It was observed that the in the lymph node of mice treated with PPP-OVA, there was a significant decrease in the frequency of Th1 cells as compared to the no treatment control. Although, there was no significant differences between no treatment control and PPP-OVA treated groups in case of activated T-helper type 1 (Th1 – CD44+Tbet+CD4+) cell frequency, there was significant decrease in this cell population in PPP group as compared to PPP-OVA group. Moreover, there was a significant increase in the activated Th2 (CD44+GATA3+CD4+) cell frequency for PPP-OVA treated group, as compared to PPP and no treatment controls. There was also a significant increase observed in proliferating Th2 (Ki67+GATA3+CD4+) cell frequency in the PPP-OVA treated group, as compared to no treatment control, however, there was an increase in this cell population in PPP group as compared to no treatment as well. As far as Treg and proliferating Treg frequencies were concerned, there was no significance found between the groups. Overall, these data suggested that in the *ex vivo* antigen recall reactions, PPP-OVA treated groups, might be generating Th2 skewing responses **(Figure 1d)**.

### PPP releasing bovine collagen type II (bc2) with a decreasing trend modulate T cell responses in collagen induced arthritis (CIA) model

To test if the decrease in RA-specific antigen exposure will modulate T cell responses in mice suffering from CIA, PPP releasing bc2 were developed. Similar to OVA, it was observed that PPP could release bc2 in a sustained and exponentially decreasing manner (**Figure 2a**). Since, it was observed that in the presence of OVA antigen, *ex vivo* cultured T cells lead to increase in the frequency of Th2 cells, we wanted to test the effect of exponentially decrease release of bc2 *in vivo* in CIA mice. Therefore, mice were induced with CIA by injection of Complete Freund’s Adjuvant and bc2 on day 0. These mice were then randomized in a cage and injected with PBS, PPP or PPP-bc2 once subcutaneously on day 14 (**Figure 2b**). Mice were then euthanized, and spleen and popliteal lymph nodes (draining lymph nodes of the paws) isolated for analysis using flow cytometry. It was observed that PPP-bc2 significantly increased regulatory T cell (Treg) frequency in the spleen, suggesting systemic suppressive responses (**Figure 2c**). Furthermore, in popliteal lymph node there was a decrease in Treg (anti-inflammatory), Th17 (pro-inflammatory), and activated and proliferating Th1 (pro-inflammatory) cell frequency in mice treated PPP bc2 compared to no treatment group. (**Figure 2c**). These data suggests that there might be an overall decrease in pro-inflammatory responses locally, since lymph nodes generally provide an indication of the responses occurring in the tissue.

**Figure 2:**
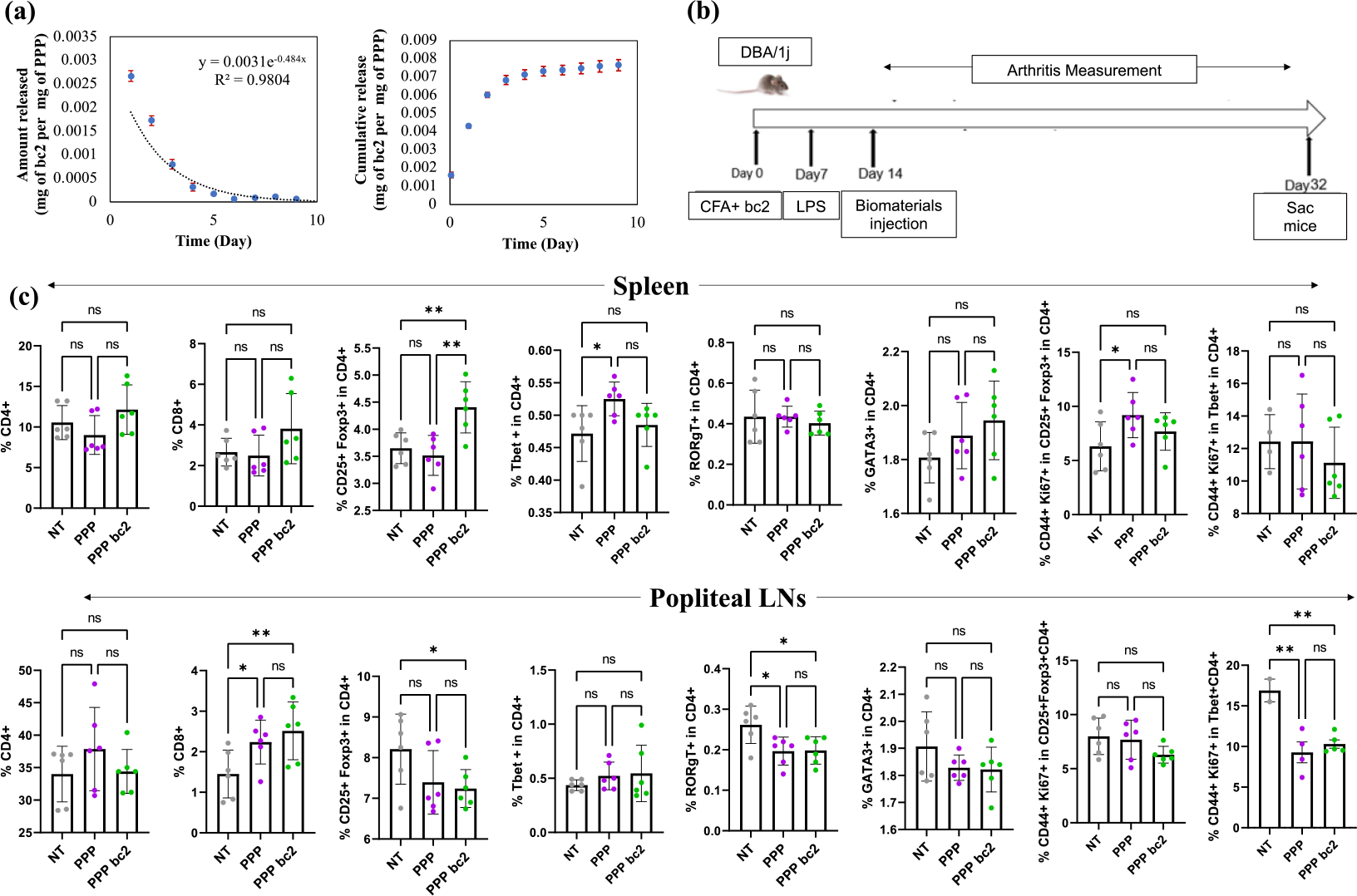
PPP releasing bc2 in an exponentially decreasing manner modulates the T cell responses in collagen induced arthritis mouse model. **(a)** PPP at 200 mg/mL release bovine collagen type II (bc2) in an exponentially decreasing manner. **(b)** Schematic of the animal experiment design. **(c)** Response of total CD4+, CD8+, regulatory T cell, Th1, Th17, Th2, activated and proliferating Treg, and activated and proliferation Th1 cells in popliteal LNs and spleen to PPP and exponentially decreasing bc2 releasing PPP-bc2. N=3 mice per group and 2 technical replicates per mice, Avg±SE, One-way ANOVA with Fisher’s LSD test, * p-value = 0.05 – 0.01, ** - p-value < 0.01, *** - p-value < 0.001.

Moreover, the effect of these formulation on physiological changes in CIA were also considered. A scoring strategy shown in **Figure 3a** was employed [15,16]. It was observed that by day 32, there was a significantly lower level of Arthritis score in PPP-bc2 treated mice as compared to PPP and no treatment mice in right rear paw (**Figure 3b**). Additionally, enzyme linked-immunosorbent assay on the serum (after dilutions, 1:5,000, 1:10,000, and 1:100,000) showed that there was not a significant change in the antibodies generated between the treatments and the controls, against mouse collagen type II, suggesting that B cells responses might not have been altered. (**Figure 3c**).

**Figure 3:**
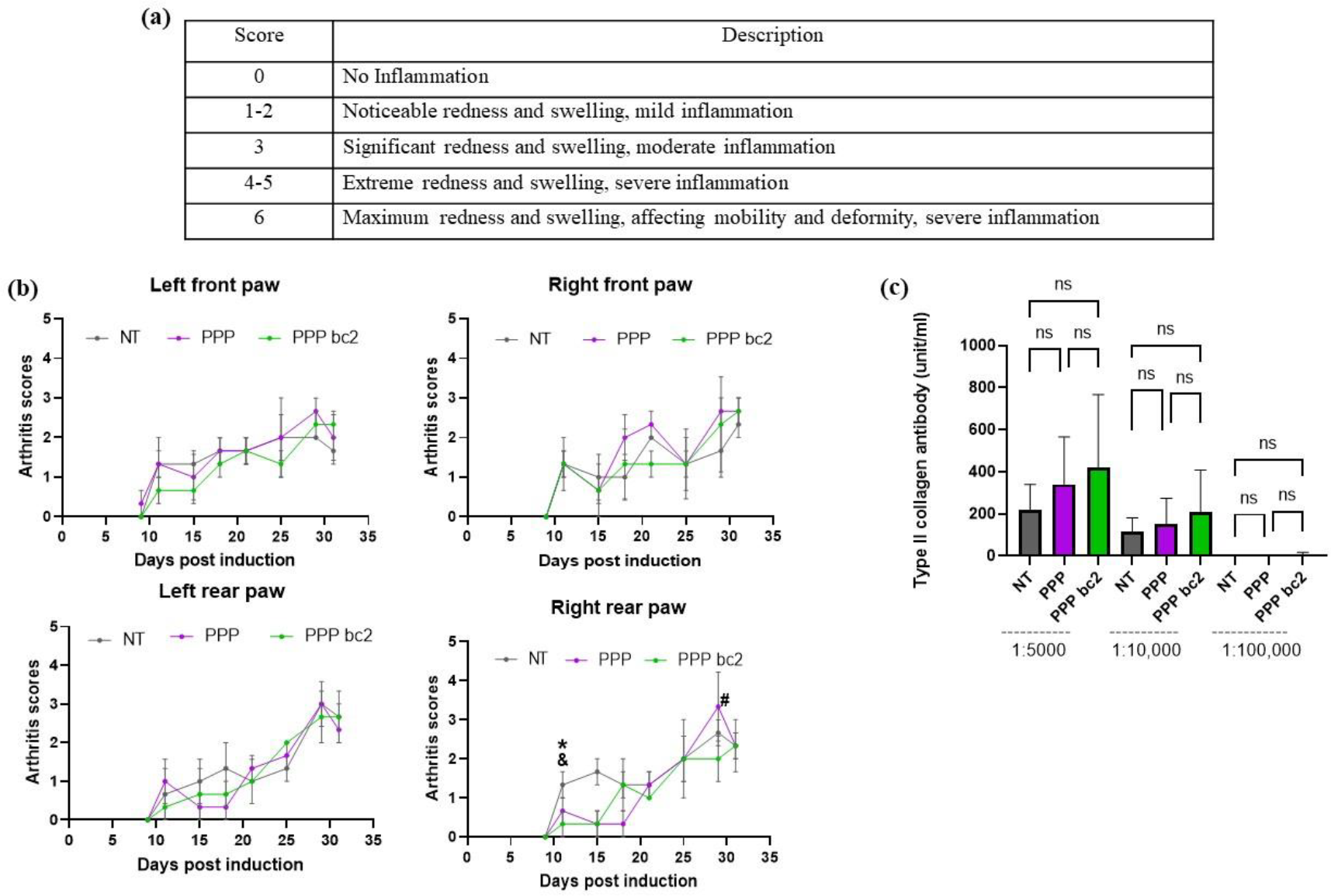
schematic of average change in RA scores of each paw and serum levels of anti-type II collagen IgG. **(a)** Scoring strategy, Paw inflammation and redness were scored on scale of 0 to 6. **(b)** Average change in RA scores of each paw from developing arthritis day to day of euthanasia, *-significant difference between PPP bc2 and NT, &-significant difference between PPP and NT, #-significant difference between PPP bc2 and PPP, * p-value = 0.05 – 0.01, N = 3 mice per group, Avg±SE, two-way ANOVA with Fisher’s LSD test **(c)** Serum collagen-specific antibody levels were assessed by ELISA, N = 3 mice per group, Avg±SE, one-way ANOVA with Fisher’s LSD test.

## Discussion

In RA that primarily affects the synovium, macrophages, CD4+ T cells, and B cells infiltrate the synovium and sometimes form a notable increase in macrophage-like and fibroblast-like synoviocytes. This condition leads to the excessive expression of enzymes, such as metalloproteinases, that can destroy the extracellular matrix and the articular structure [17–22]. To understand the mechanism of the disease, active (antigen-induced arthritis) induction was chosen in this study. To understand the role of temporal changes in the concentration of antigen in the CIA model, an exponentially decreasing dose pattern of bc2 was chosen.

Th1 and Th17 cells are key participants in RA. For example, Th1 via expression of cytokine can activate macrophages and enhance the ability of antigen-presenting to the naïve T cells. Furthermore, pro-inflammatory cytokines released by Th1 can induce synovial inflammation, recruit more immune cells, and contribute to joint destruction. In addition, Th1 can indirectly contribute to bone erosion by promoting osteoclastogenesis. Th17 by producing IL-17 can stimulate fibroblast-like synoviocytes and also have a role in bone erosion and recruitment of neutrophils [23–25]. The results (**Figure 2c)** indicated that this reduced antigen availability leads to a decline in the Th17 in PPP bc2 treated mice compared to no treatment mice and activated and proliferated Th1 cell population in PPP bc2 treated mice compared to no treated mice in secondary lymphoid organs (popliteal LNs)

An increase in the number of Tregs is helpful in the treatment of RA [15,26–28]. Treg by maintaining immune balance, production of cytokines, regulation of antigen-presenting cells, and direct cell-cell contact suppression leads to apoptosis of effector T cells and immunosuppression [29–31]. Our finding (**Figure 2c)** suggested that the exponentially reduced dose regime of bc2 leads to a considerable increase in the Treg cell population in PPP bc2-treated mice compared to untreated mice in the secondary lymphoid organ (spleen).

Autoantibody production by B cells is a marker of diagnostic and prognostic in RA [32]. However, our results (**Figure 3c**) showed there is no significant reduction in collagen-specific immunoglobulin (IgG) in the serum of PPP bc2-treated mice compared to other groups. These data suggested that a more potent antigen specific response might be required for generating a viable therapy against autoimmune RA, which is a limitation of this study. Nonetheless, this study provides an insight in the idea of antigen release profiles affecting adaptive immune outputs.

Last but not least, in wild-type C57BL/6j mouse model, it was observed that in mice treated with the same adjuvant (PPP) and different antigen (OVA) there is a significant decrease in the frequency of Th1 cells as compared to the no treatment control and a significant increase in the activated Th2 in the *ex vivo* **(Figure 1d)**.

## Experimental section

### Materials and Methods

#### Thermoresponsive drug delivery system

PLGA-PEG-PLGA (1000-1000-1000) (PPP, Sigma Aldrich) were purchased and dissolved in 1X phosphate buffered saline at 4 °C overnight by gentle vortexing at 200 mg/mL concentration. OVA or BC2 were weighed and added to the PPP solution and mixed well at 4 °C. This mixture was used immediately.

#### Release kinetics

Release kinetics of OVA or BC2 was determined by incubating 10 mg of the PPP containing the protein or without the protein in 1 mL of PBS in centrifuge tubes. Triplicates of each sample were placed on in a bead-bath (Thermo Fisher Scientific) at 37 °C. Next, 800 μL of the supernatant was carefully transferred to a new 1.5 mL centrifuge tube (Eppendorf, Hauppauge, NY) and stored in -80 °C until use. Then 800 μl was replaced with new buffer and the tube was then placed in the bead bath. This was repeated for 9 days. The amount of protein released by PPP was then quantified using BCA assay (Thermo Fisher Scientific).

#### OVA treatment model

All experimental procedures were conducted in accordance with the guidelines and approval of the Institutional Animal Care and Use Committee (IACUC) at Arizona State University, protocol number 19-1712R for these experiments. C57BL/6j mice, aged 6-8 weeks were obtained from Jackson Laboratories. Mice were injected with PPP, PBS or PPP loaded with OVA on day 0 subcutaneously, in 50 μL of PBS. Next, the mice were sacrificed on day 7, and spleen and inguinal lymph nodes were isolated for analysis using flow cytometry.

#### CIA induction in DBA/1j mice

All experimental procedures were conducted in accordance with the guidelines and approval of the Institutional Animal Care and Use Committee (IACUC) at Arizona State University, protocol number 19-1712R for these experiments. DBA/1j mice, aged ten to twelve weeks, were obtained from Jackson Laboratories. To induce rheumatoid arthritis (RA), on day 0, a subcutaneous injection was administered at the base of the tail, consisting of a 1:1 volume ratio mixture of Complete Freund’s adjuvant (prepared from Incomplete Freund’s adjuvant and Mtb) (Difco via Fisher, IFA cat# DF0639-60-6, Sparks, MD & Mtb H37Ra cat# DF3114-33-8, Sparks, MD) along with bc2 antigen (Chrondrex via Fisher, Cat # 50-152-6983, Woodinville, WA). Mice were also injected with PPP or PPP-bc2 on day 14, and the mice were then sacrificed on day 32, and spleen, blood and popliteal lymph nodes were isolated for flow cytometry analysis.

#### CIA scores and measurements

The progression of CIA symptoms was closely monitored and assessed three times per week. Baseline measurements of paw size and the mice’s weights were recorded for reference. Paw inflammation was evaluated using a scoring system ranging from 0 to 6, with scores of 3 indicating moderate inflammation and scores higher than 3 indicating severe inflammation. Additionally, paw measurements were taken using vernier calipers to track any changes in size. To ensure the well-being of the mice, their weights were monitored three times a week, and they were provided with wet food (food pellets mixed with water) to facilitate easy access to nutrition.

#### Ex vivo culture of T cells

After the completion of animal studies, the spleens from the mice were extracted for recall reaction experiments. Plates were coated with anti-CD3 (0.5 μg/mL) and incubated at 37 °C for 2 hours. The isolated splenocytes were then incubated in 3 mL of 1x red blood cell lysis buffer at 4 °C for 5 minutes. After centrifugation at 300x g for 5 minutes and discarding the supernatant, the splenocytes were resuspended in RPMI-1640 Medium with L-glutamine (VWR, Radnor, PA), 10% fetal bovine serum, and 1% penicillin-streptomycin (VWR, Radnor, PA). The plates were washed twice with 1x PBS to remove the anti-CD3 coating. A total of 1x10^6 splenocytes/mL were seeded in the presence of OVA or BSA (10 μg/mL). For splenocyte stimulation, anti-CD28 (0.5 μg/mL) and IL-2 (10 ng/mL) were added. Antibody staining and flow cytometry analysis were performed 72 hours after culturing the splenocytes at 37 °C.

#### Flow cytometry

Flow cytometry analysis was conducted using the Attune NXT system (BD Biosciences, BioLegend, Invitrogen, ThermoFisher, Waltham, MA, USA). On day 100, cells from the inguinal lymph nodes, popliteal lymph nodes, and spleen of CIA mice were collected. These cells were then subjected to staining with specific fluorescent antibodies in a 1% Fluorescent Activated Cell Sorting (FACS) buffer, prepared by combining 0.001% bovine serum albumin, 0.5M EDTA, and 0.0001% NaN3 in 1xPBS. The following antibodies were utilized for the experimental analyses.

| Target | Fluorophore | Detector | Company | Catalog number | Clone |
| --- | --- | --- | --- | --- | --- |
| CD4 | PE | YL1 | BD Biosciences | 12-0041-82 | GK1.5 |
| CD8 | APC-R700 | RL2 | BD Biosciences | 564983 | 53-6.7 |
| CD25 | PECy7 | YL3 | BD Biosciences | 552880 | PC61 |
| Tbet | BV785 | VL6 | BioLegend | 644835 | 4B10 |
| FoxP3 | eF450 | VL1 | Invitrogen | 48-5773-82 | FJK-16s |
| RORγT | BV650 | VL4 | BD Biosciences | 564722 | Q31-378 |
| GATA3 | BV711 | VL5 | BD Biosciences | 565449 | L50-823 |
| CD44 | BV480 | VL2 | BD Biosciences | 566200 | IM7 |
| Ki67 | FITC | BL1 | Invitrogen | 11-5698-82 | SolA15 |

#### Enzyme Linked Immunosorbent Assay

Microwells were coated with 100 μL of Coating Buffer containing 20 μg/mL bc2 per well. For the standard wells, microwells were coated with unlabeled mouse IgG1 Next, the plate was incubated overnight at 4°C. The wells were then aspirated and washed three times with approximately 300 μL of Wash Buffer per well. The plates were then blocked with 200 μL of 1% BSA per well and incubated at room temperature (RT) for at least 1 hour. The wells were then aspirated and washed once. Samples in assay Diluent were generated, with serum sample dilution ranges of 1:500, to 1:100,000. A 100 μL of each sample and control were added into their respective wells, and sealed for 1 hour incubation at RT. The wells were then aspirated and a total of 5 washes were performed. Next, 100 μL of prepared Detection Antibody diluted in Assay Diluent to each well (at 1:5000 dilution) was added, and the plate was sealed for 1 hour incubation at RT. The wells were then washed 5 times, and 100 μL of prepared anti-mouse IgG/HRP diluted in Assay Diluent to each well (dilution: 1:1000) was added, and sealed for 30 minutes incubation at RT. Next, the wells were washed 7 times, allowing the wells to soak in wash buffer for 1 minute for each wash. Next, 100 μL of Substrate Solution was added to each well and incubated t at RT in the dark. Finally, 20 μL of Stop Solution was added to each well, and the absorbance was read using a plate reader at 450 nm within 30 minutes of stopping the reaction.

#### Statistical Analysis

The data are presented as mean ± standard error. Statistical comparisons among multiple treatment groups were conducted using two-way and one-way analysis of variance (ANOVA) with Fisher’s least significant difference (LSD) test. A p-value of less than 0.05 was considered statistically significant. GraphPad Prism Software 6.0 (San Diego, CA) was used for the statistical analysis.

## Conclusion

Our findings shed light on the potential of using PPP with a decreasing trend in OVA release as a strategy to modulate T cell responses and provide insights into the regulation of immune responses. Moreover, this study also provides for the first time a drug delivery strategy to control T cell responses only based on antigen release in vivo.

## Authors’ contributions

Arezoo Esrafili designed and performed the experiments, analyzed data and wrote the manuscript. Other authors performed experiments. Abhinav P. Acharya and Mattew Peet helped design experiments, procured funding and wrote the manuscript.

## Acknowledgement

The authors acknowledge the support of 1R01GM144966-01 from National Institute of Health, US to APA and MMP.

