## Supplementary Figure 1 and 2 for "Exponentially decreasing exposure of antigen generates anti-inflammatory T-cell responses"

### Supplementary Figures

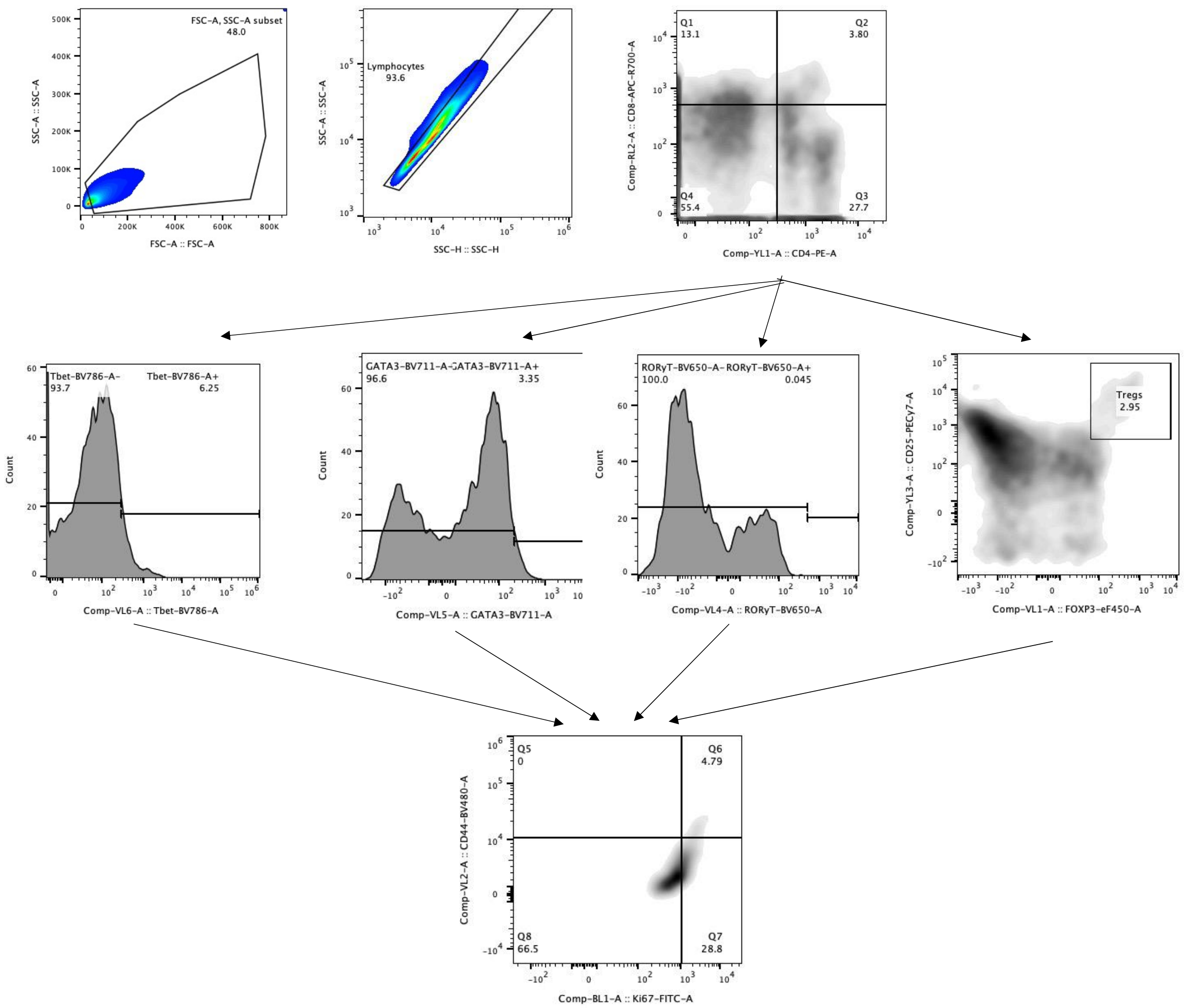


**Supplementary Figure 1: Flow cytometry schematic for the analysis of the immune cells for both *in vivo* and *ex vivo* analysis.**

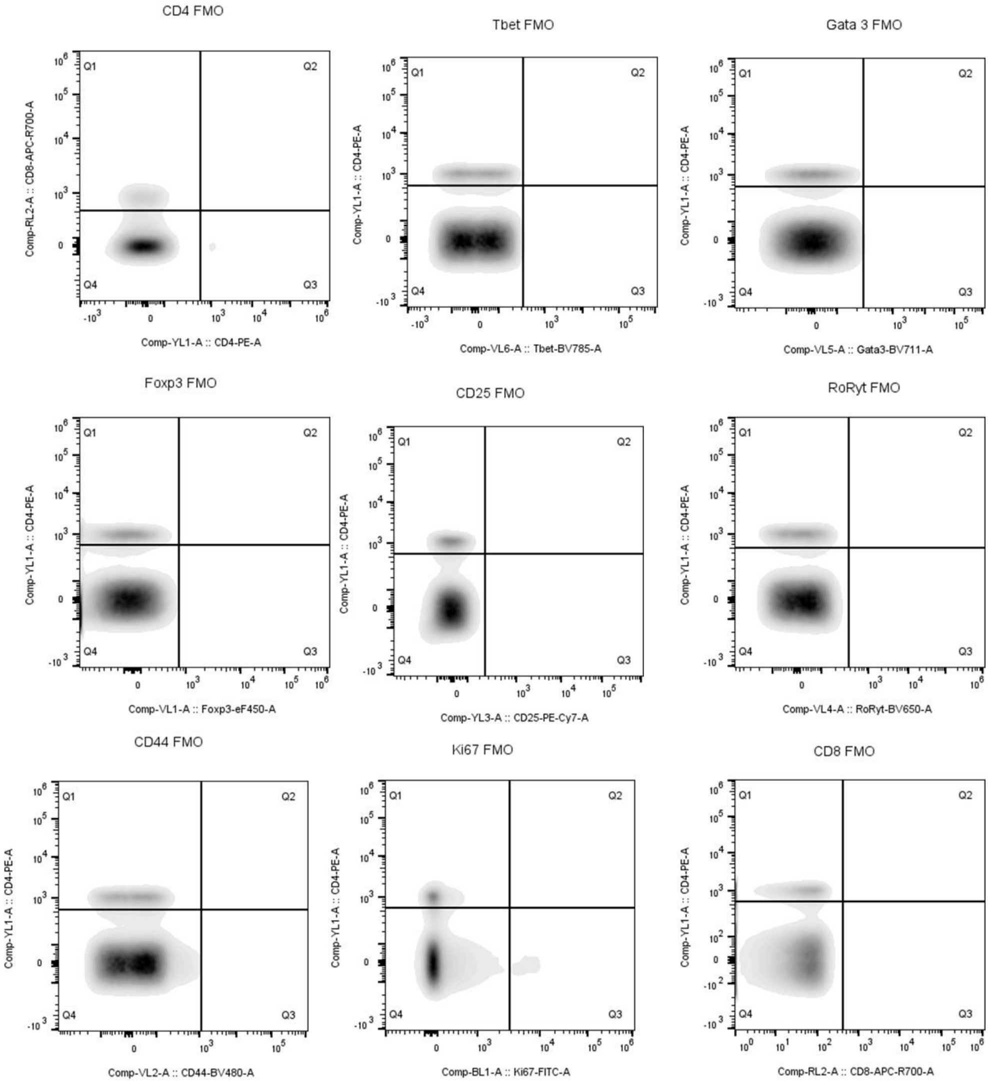


**Supplementary Figure 2: Flow cytometry fluorescence minus one (FMO) to determine the negative controls for each of the fluorophore utilized.**
